# Automated Detection of Quiet and Non-Quiet Sleep in Preterm Neonates from aEEG: Towards Predicting Brain Maturation

**DOI:** 10.64898/2026.02.09.704893

**Authors:** Théophile De Backer, Albert Fabregat-Sanjuan, Jordi Solé-Casals, Vicenç Pascual-Rubio, Rosa Pàmies-Vilà

## Abstract

**Background:** Preterm birth is associated with an increased risk for neurodevelopmental impairments, requiring brain monitoring using amplitude-integrated electroencephalography (aEEG). While established tools detect severe dysfunction (e.g., Hellström-Westas classification), methods for assessing mild to moderate impairments—such as Burdjalov scoring or expert-based Sleep–Wake Cycle identification—are subjective and require specialized training. Automated neonatal sleep-staging models usually rely on polysomnography from term infants, a resource-intensive method rarely feasible in NICUs, where simplified single-channel aEEG is standard.

**Methods:** aEEG recordings from 40 neurologically healthy neonates (32–42 weeks PMA) were collected and annotated for quiet (QS) and non-quiet sleep (NQS) by an expert clinician. Signals were bandpass filtered, segmented into 30 s epochs, and cleaned using impedance thresholds. 69 temporal, spectral, wavelet, EMG-inspired, and aEEG-envelope features were extracted. The 5 most relevant features were selected for QS/NQS classification using several machine-learning models validated with leave-one-subject-out cross-validation. A partial least squares model was then trained on QS-derived features to predict postmenstrual age and assess correlations with brain maturation.

**Results:** The k-Nearest Neighbors (KNN) classifier showed the best QS/NQS discrimination, with mean Cohen’s *κ* = 0.69 ± 0.14 for preterm (32–37 weeks PMA) and 0.48 ± 0.21 for term infants. QS-derived features correlated strongly with postmenstrual age (PMA). The PLS model predicted PMA with an average error of 0.88 weeks (MSE = 1.33 weeks, r = 0.91), while the fully automated version using predicted QS segments yielded an error of 1.08 weeks (r = 0.86).

**Conclusion:** Automated QS/NQS detection from single-channel aEEG is feasible in preterm neonates. Despite reduced accuracy in term infants, QS-derived features closely track brain maturation, supporting the potential of aEEG-based models for objective, early detection of neuromaturation delays in preterm infants

## Introduction

Each year, nearly one in ten neonates is born preterm (before 37 weeks of gestational age), according to the World Health Organization (WHO) [1]. Preterm birth is associated with an increased risk of neurodevelopmental conditions such as intraventricular hemorrhage (IVH) [2], cerebral palsy [3], autism spectrum disorder, and attention-deficit/hyperactivity disorder (ADHD) [4, 5]. Consequently, Neonatal Intensive Care Units (NICUs) have developed specialized techniques to monitor early brain development in this vulnerable population.

One of the most widely used techniques is amplitude-integrated electroencephalography (aEEG), a simplified version of conventional EEG designed for continuous bedside monitoring of neonates which allows easy recognition of its patterns by non-expert personnel in clinical neurophysiology. Unlike standard EEG used during polysomnography (PSG), which typically requires between 8 and 18 electrodes [6], aEEG requires only five. This reduction makes it more suitable for the smaller head size of neonates [7], causes minimal sleep disturbance [8], and makes it more practical for daily clinical use in NICUs [9]. The electrodes are placed according to the international 10–20 system [10], typically with two on each hemisphere (e.g., P3–P4 or C3–C4) and one at the vertex (Cz) serving as ground or reference, thus providing one channel per hemisphere. The raw EEG signals are then transformed into aEEG traces following a specific processing pipeline [11]. For improved clinical interpretation, these traces are displayed on a semi-logarithmic scale—voltages below 10 µV are plotted linearly, while higher voltages are shown logarithmically—and visualized over 4-hour windows (6 cm/h).

aEEG has proven highly effective for detecting severe abnormalities, such as seizures [12, 13] or encephalopathy patterns defined by Hellström-Westas [14]. However, its ability to identify subtler abnormalities—such as mild brain dysfunction, which are interpreted as delayed neuromaturation—remains limited [15]. To assess brain maturation, clinicians often use the Burdjalov scoring system [16], which defines typical aEEG features at different postmenstrual ages (PMAs). By comparing the maturation age estimated from the Burdjalov score with the neonate’s actual PMA (based on gestational age at birth and postnatal age), clinicians can evaluate whether brain maturation is appropriate or delayed.

Despite its widespread use, this scoring method depends heavily on subjective visual interpretation and has shown limited inter-rater reliability [17]. In practice, clinicians instead prefer to evaluate Sleep–Wake Cycles (SWC)—periods of alternating quiet sleep (QS) and non–quiet sleep (NQS)—as their presence and duration are strong indicators of brain maturation [18, 19]. Nevertheless, manually labeling SWC on long aEEG recordings is labor-intensive, subjective [20], and requires expert training [15].

In PSG studies, clinicians can manually classify four distinct sleep stages: wakefulness, active sleep (AS), quiet sleep (QS), and intermediate sleep (IS), which combines characteristics of QS and AS [21, 22]. This fine-grained classification is possible because PSG recordings typically include multiple EEG channels (from 9 to over 18 electrodes) and complementary physiological signals such as electro-oculogram (EOG) and electrocardiogram (ECG) [23]. Data are reviewed in 30-second intervals, as recommended by the American Academy of Sleep Medicine (AASM) [24]. EEG channels help identify patterns such as the tracé alternant—linked to QS—while EOG and ECG confirm additional features (e.g., rapid eye movements for AS, lower heart rate for QS) [25]. Although this detailed approach allows precise identification of sleep stages, it is extremely time-consuming [26] and requires complex setups. Therefore, PSG is rarely used in preterm infants outside of research settings [9, 27], since the application of multiple electrodes can cause skin injury or infection in these fragile patients [28].

aEEG was developed precisely to overcome these limitations. With its simple five-electrode setup and capacity for long-term recordings (up to 24 hours) at the bedside, it enables continuous monitoring with minimal disruption to the infant’s sleep. Although the reduced number of channels prevents the detailed four-stage classification possible with PSG, aEEG data allow a reliable two-stage distinction—quiet sleep (QS) versus non–quiet sleep (NQS). According to Zhang et al. [11], alternating periods of quiet sleep (QS) and non-quiet sleep (NQS) can be identified in aEEG recordings, forming Sleep–Wake Cycles (SWC) that are typically analyzed over 4-hour windows rather than the 30-second epochs used in polysomnography (PSG).

Most previous research on neonatal sleep staging has relied on PSG data, proposing models for two-, three-, or four-stage classification [29–39]. Some models use up to eight EEG channels [29, 36, 37], but their performance decreases significantly when fewer channels are available, as demonstrated by Ansari et al. [40] and Montazeri [41]. In one of the earliest demonstrations of automated neonatal sleep staging, Koolen et al. [42] developed a classifier capable of distinguishing sleep states from multi-channel EEG across different postmenstrual ages, highlighting both the feasibility and the challenges of automating neonatal sleep assessment.

Consequently, recent efforts have focused on single-channel models to enable sleep staging with EEG data compatible with aEEG systems. Most studies simulated this condition by selecting a single channel from multi-channel PSG datasets [35, 40, 43, 44], while only Montazeri et al. [41] used EEG data recorded under aEEG standards—though exclusively from term neonates (± 40 weeks PMA).

Among existing QS/NQS classification models, Ansari et al. [40] trained a sinc-block convolutional neural network achieving a Cohen’s Kappa of 0.75, while Montazeri et al. [41] obtained 83% accuracy and Kappa = 0.71 using a convolutional neural network validated on external term data. Irfan et al. [35] achieved 87% accuracy and Kappa = 0.74 using an ensemble model with extensive feature extraction (147 features). However, none of these studies validated their models on preterm infants (*<* 37 weeks PMA) using data recorded under aEEG standards.

Based on these clinical and methodological gaps, the present study pursues two objectives. The first objective is to develop a machine learning model capable of automatically classifying quiet sleep (QS) and non–quiet sleep (NQS) periods from long-term neonatal EEG recordings acquired under aEEG standards. The model is designed to operate with a maximum of two EEG channels, ensuring compatibility with the simplified aEEG setup commonly used in NICUs. Furthermore, it should be robust across a wide range of postmenstrual ages (32–42 weeks in our work) and capable of detecting relatively infrequent QS segments (approximately 20% of the data) within recordings dominated by NQS periods and potential artifacts.

The second objective is to explore the automatic estimation of postmenstrual age (PMA) based on features extracted from neonatal EEG and aEEG signals. Specifically, the study investigates whether features derived exclusively from QS segments provide a stronger correlation with PMA than those obtained from unfiltered datasets containing both QS and NQS segments. To this end, two models are compared: one trained solely on QS data, either manually labeled or automatically detected, and another trained on the original class distribution (80% NQS, 20% QS).

Accordingly, this study addressed two key research questions: (RQ1) Can a machine learning model be developed for classifying QS and NQS segments in neonates, using no more than two channels of long-term EEG recordings across a broad range of postmenstrual ages (32–42 weeks)? and (RQ2) How strongly do features extracted from QS segments correlate with PMA, and does filtering recordings to retain only QS segments improve PMA prediction compared with using unfiltered data?

## Materials and methods

### Dataset

Forty neurologically healthy neonates were recruited from Hospital Universitari Sant Joan de Reus (HUSJR), Tarragona, Spain. This work was performed in the framework of a clinical trial conducted at HUSJR, which was approved by the Ethics Committee of the Institut d’Investigació Sanitària Pere Virgili (IISPV) with reference numbers 032/2021 and 127/2024, and authorized by the Spanish Agency of Medicines and Medical Devices (AEMPS) with reference 918/21/EC-R. [45]. Written informed consent was obtained from the parents or legal guardians of all participants prior to their inclusion in the study.

In order to investigate the correlation between the aEEG data and the brain age maturation, it was crucial that all the neonates recruited were neurologically healthy such that their aEEG signal would not be divergent from what is considered normal at their PMA. For this, the inclusion criteria for the neonates were that they had no signs of encephalopathies, were less than five days old at the time of the recording, and had a minimum score of five on the Apgar test taken at five minutes of life. Also, the first blood pH value was taken into consideration, with a minimum required value of 7.2. Alterations in cerebral echography were also monitored to ensure the neonates were neurologically healthy.

In Table 1, descriptive statistics regarding the PMA of the neonates and the recording length are provided. The distribution of neonates per PMA category is visible in Figure 1. In total, the dataset included more than 1100 hours of aEEG recording, with a mean recording length of 27 hours per neonate.

**Table 1.** Dataset PMA and Recording Duration.

| Metric | Value |
| --- | --- |
| Mean PMA (weeks) | 36.71 |
| Standard Deviation of PMA | 2.45 |
| Minimum PMA (weeks) | 32.57 |
| Maximum PMA (weeks) | 41.43 |
| Mean Recording Duration per subject (hours) | 27.73 |
| Total Recording Hours | 1109.19 |
| Minimum Recording Duration (hours) | 16.62 |
| Maximum Recording Duration (hours) | 47.20 |

**Fig 1.**
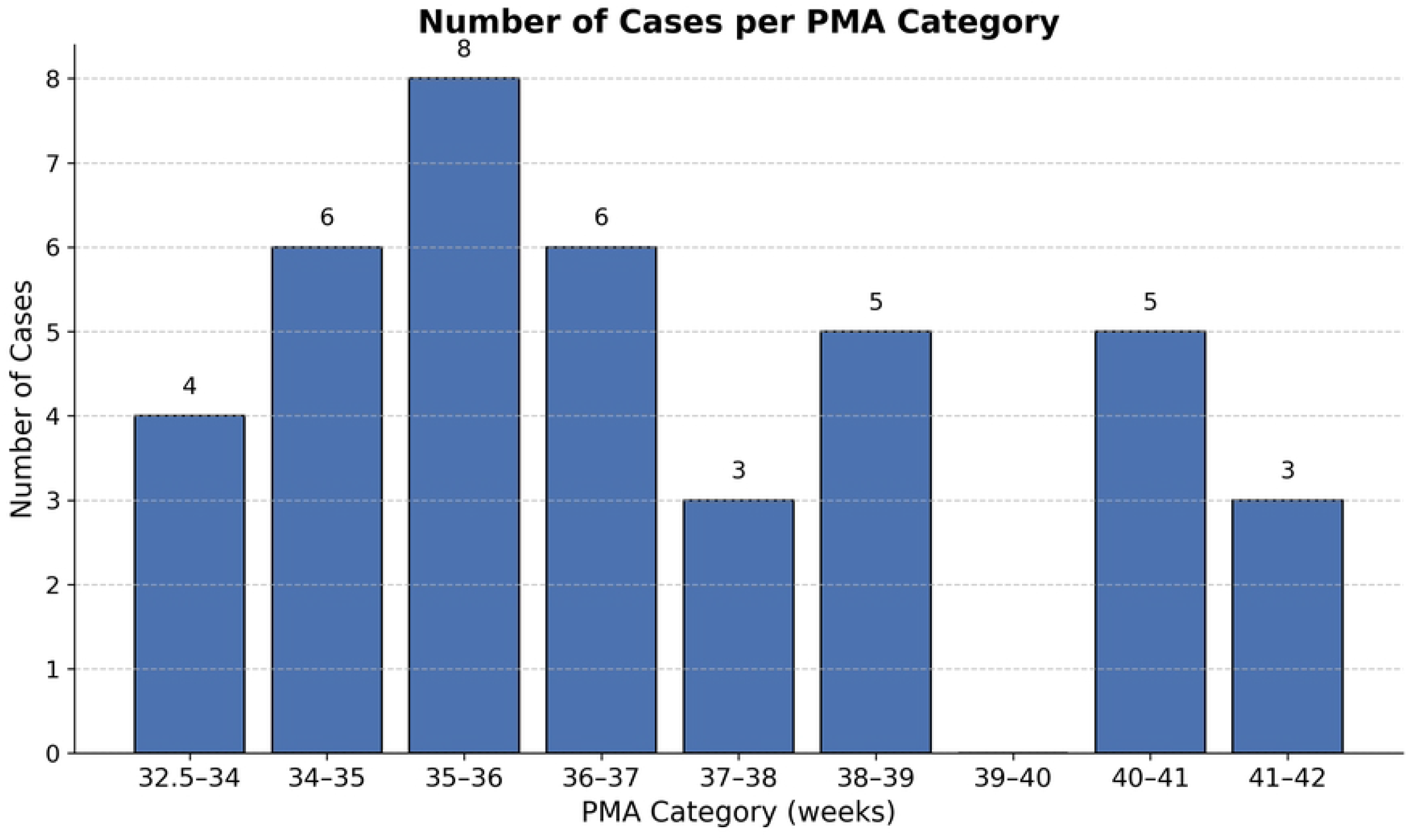
Histogram showing the number of available neonates in each PMA category

Each recording was performed using a five-electrode montage with C3, C4, P3, P4, and Cz as the ground electrode, according to the 10-20 international convention. C3 and P3 were used to monitor the left hemisphere, while C4 and P4 monitored the right hemisphere. All electrodes operated at a sampling frequency of 200 Hz with impedance monitored at a sampling rate of 100 Hz.

All data from the 40 neonates were visually labeled for sleep-wake cycles (periods of QS) by the same expert clinician. For this task, the clinician had access to the aEEG, EEG, and impedance signals from one hemisphere at a time. Representative examples of amplitude-integrated EEG (aEEG) recordings illustrating the annotation and envelope extraction process are shown in Figures S1 and S2. No other polysomnography channels such as electrooculography (EOG), electromyography (EMG) or electrocardiogram (ECG) were available to the clinician during the annotation.

### Pre-processing

Although both hemispheres were labeled, we used the hemisphere with the lower electrode impedance. The pre-processing pipeline then consisted of three parts: bandpass filtering, signal segmentation, and impedance filtering (see Figure 2).

**Fig 2.**
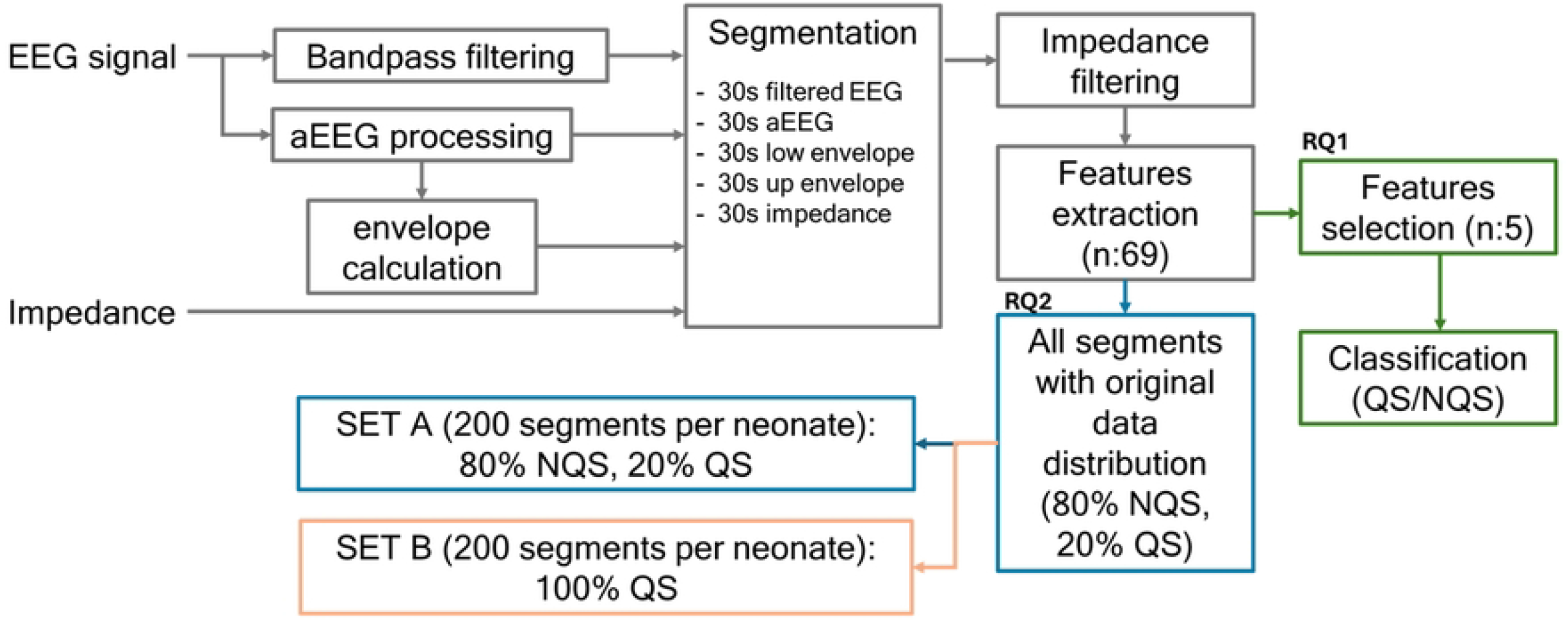
The pre-processing pipeline is indicated by the grey blocks. RQ1 includes a feature selection step, keeping only the top five features for training the classifiers. RQ2 uses all 69 extracted features to create two sets. Set A contains 200 segments per neonate, following the original class distribution, while set B is composed of exclusively QS segments.

The raw EEG signals, obtained in European Data Format (EDF) files from the hospital monitor, were first filtered using a bandpass filter of 0.3–35 Hz. This is widely used in the literature and was chosen because high and low frequency elements are often associated with noise and artifacts; thereby, filtering them out improves the signal-to-noise ratio. The aEEG signal and envelope of the aEEG were instead processed from the raw EEG signal following their respective processing algorithm detailed in [11, 46] .

Then the signal was segmented in 30 s non-overlapping segments. This duration is the most commonly used for studies on neonatal sleep-staging using EEG signals and was used both in [35] and [40]. For each segment, the EEG, aEEG, and impedance data points, as well as data points corresponding to the upper and lower envelopes of the aEEG signal, were kept. Since the impedance was recorded at half the EEG sampling frequency (200 Hz), and the upper and lower envelopes contained fewer data points due to the smoothing window described in [11, 46], interpolation was applied.

Finally, the label (0: NQS or 1: QS) assigned by the clinician that corresponded to this 30 s segment was stored.

The final part of the pre-processing pipeline is impedance filtering. At 20 kΩ, the electrode saturates, preventing an accurate reading of the impedance value. For this reason, all segments where one or both electrodes exceeded this threshold were removed. Occasionally, impedance peaks occurred sporadically. Since they can affect the beginning and end of sleep-wake cycle patterns, making it difficult for clinicians to accurately annotate quiet sleep periods, a second impedance filtering was applied using a 30-minute sliding window over the entire recording. Through iterative testing, it was determined that removing windows containing 10 or more peaks within a 30-minute period—or 10 peaks per 60 segments—effectively discarded data segments affected by high impedance during labeling while retaining most (77%) of the data.

### Feature extraction

A total of 69 features were extracted from each 30-second EEG segment. Temporal features, such as mean amplitude, variance, and peak-to-peak amplitude, were computed to characterize the basic morphology of the EEG signal. Frequency-domain features, including log-transformed band powers, spectral entropy, and spectral edge frequency, were calculated to describe its spectral composition and complexity. DWT-based features were derived to capture multi-scale temporal variations in the EEG according to [35]. To broaden the analysis beyond conventional neonatal sleep-staging literature, features commonly used in EMG signal analysis were also incorporated to assess their potential for distinguishing between QS and NQS. These included the Log Detector (LOG) and Myopulse Rate (MYO), both computed for each EEG segment. Finally, aEEG- and envelope-derived features were extracted, following Yasova Barbeau and Weiss [27], who reported that the difference in bandwidth of the aEEG trend—defined as the amplitude gap between the upper and lower envelopes—is an important clinical indicator for identifying QS periods. In Table 2, the most relevant features are defined, while the complete set of extracted features is provided in Table S1.

**Table 2.** Definition of most relevant selected features: DWT-based, spectral, EMG-inspired, and aEEG/envelope features. When not explicitly specified on aEEG, the features are computed using the 30s EEG signal segment.

| Feature | Description |
| --- | --- |
| **DWT-based features** |  |
| D3_AbsMean_Ratio | Ratio of absolute mean values between adjacent DWT sub-bands (D3/D2) |
| D4_AbsMean_Ratio | Ratio of absolute mean values between adjacent DWT sub-bands (D4/D3) |
| **Spectral features** |  |
| Spectral entropy | Shannon entropy of the normalized power spectral density |
| Log spectral edge | $\log(\text{frequency below which a fixed percentage of power resides, e.g., 90\%})$ |
| **EMG-inspired features** |  |
| Log detector (LOG) | $\exp\left(\frac{1}{N} \sum_{i=1}^N \log(\vert X_i\vert )\right)$: sensitive to low-level activity |
| Myopulse rate (MYO) | Percent of time the signal exceeds $2\times$ its mean amplitude |
| **aEEG and envelope features** |  |
| Mean lower envelope | Average amplitude of the aEEG lower envelope |
| Envelope difference | Difference between upper and lower aEEG envelopes |
| aEEG_spectral_flatness | Ratio of geometric to arithmetic mean of aEEG power spectrum |

### Feature selection/reduction

A step unique to RQ1 was the elimination of redundant features for the QS/NQS classification task (see Figure 2). For this, it was decided to use the Gram-Schmidt orthogonal feature selection algorithm as it returns an ordered list of features ranked by their correlation with the output while maximizing linear independence among the selected features [47].

Also, it was decided to keep only the five strongest features in order to develop a model that maintains a certain level of interpretability. In fact, having a lot of features makes it difficult to predict how a model will behave given a certain input [48].

Because the Gram-Schmidt algorithm normalizes features and labels, we additionally balanced the dataset through under-sampling to ensure equal numbers of QS and NQS segments were presented. This prevented the mean from being biased toward the majority (NQS) class.

Finally, to improve the robustness of the results, the algorithm was run 10 times with different random seeds for selecting balanced samples. The key steps of the algorithm are summarized in Algorithm 1.

### Classification methods

For classification, five traditional machine learning models including Linear Discriminant Analysis (LDA), k-Nearest Neighbors (KNN), support vector machine (SVM) with a linear kernel, decision tree (DT), and random forest (RF) were trained using default hyperparameters. The exception was the KNN, where the k-value was set to 50 due to the large dataset size. Additionally, to prevent overfitting and reduce computational expense, the maximum depth for both the DT and RF models was limited to 10.

#### Algorithm 1

Gram-Schmidt Feature Selection

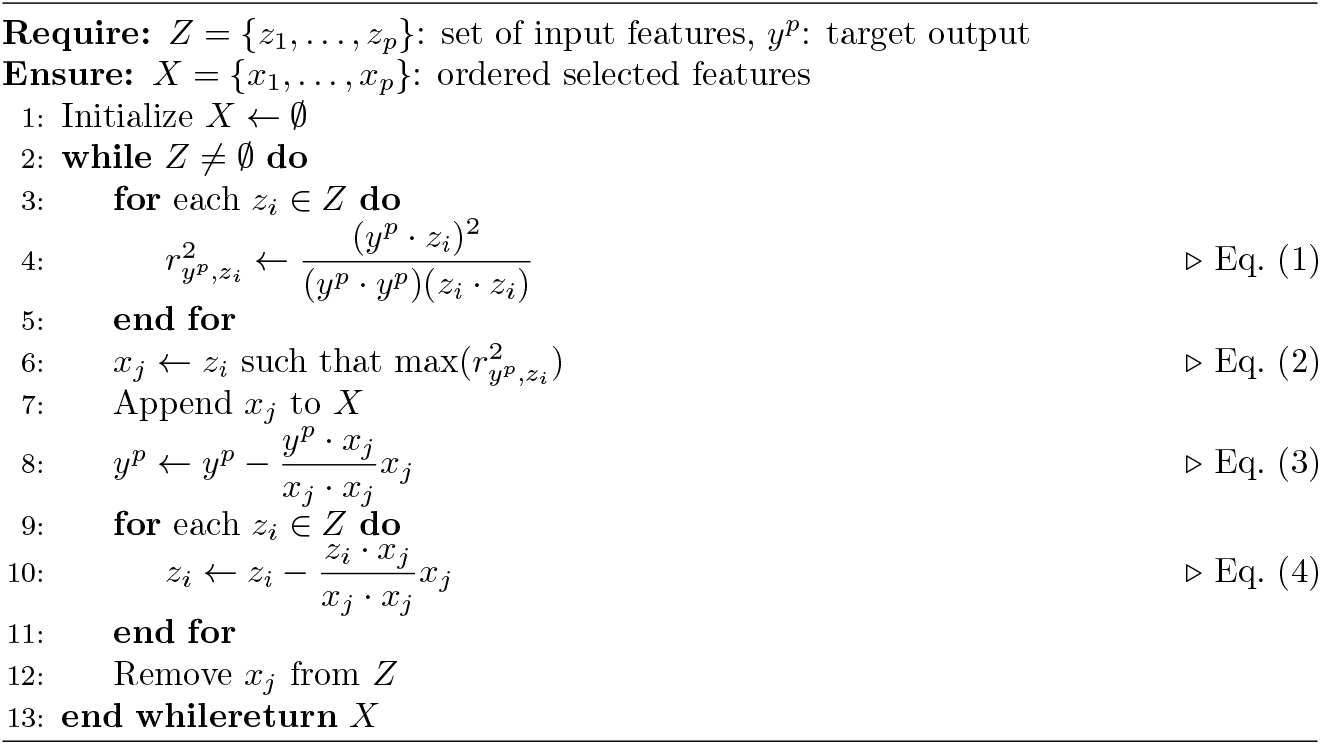

For the choice between Leave-One-Subject-Out (LOSO) and k-fold cross-validation, both commonly used in the literature, we designed an experiment to test whether each subject’s data clusters closely around them. If so, this would justify using subject-level validation (LOSO), as opposed to k-fold cross-validation, which does not guarantee subject-level independence and may include data from the same subject in both training and validation sets.

First, two cases with identical PMA were randomly selected from the dataset. Each case was segmented and pre-processed as described in the previous sections, ensuring that high-impedance segments were removed from both cases. Feature extraction was performed on each case, and the top five features identified using the Gram-Schmidt algorithm were retained to represent the data in five dimensions. Each case was then resampled to match the length of the shortest one, ensuring equal length for comparison. Finally, the inter- and intra-subject distances were calculated using the Euclidean distance between feature vectors and compared using a t-test. The results of this small experiment are presented in Table 3 demonstrating that even for two neonates of equal PMA, the intra-subject distance of each neonate is smaller than the inter-subject distance. This suggests that datapoints corresponding to one neonate are closely distributed to each other. The t-test confirmed that the inter-subject and combined intra-subject distance groups were statistically different from each other.

**Table 3.** Mean Euclidean Distance Comparison.

| Subject | Comparison | Mean Distance |
| --- | --- | --- |
| Subject 1 | Intra-subject | 6.4811 |
| Subject 2 | Intra-subject | 1.5293 |
| Subject 1 and 2 | Inter-subject | 9.0976 |
| **T-test (Intra vs Inter):** $t = -697.7842$, $p = 0.0000$ | | |

The five trained classifiers were thus validated using Leave-One-Subject-Out (LOSO) cross-validation to preserve case independence. The evaluation metrics included accuracy, precision, recall, F1-score, and Cohen’s Kappa. Definitions for each of these metrics are provided below:

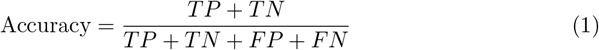

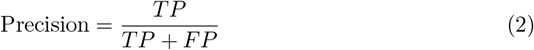

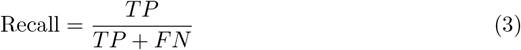

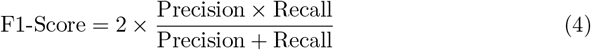

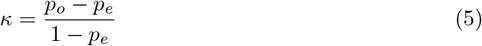

where TP, TN, FP, and FN denote the true positive, true negative, false positive, and false negative values of the confusion matrix, respectively. *κ* represents Cohen’s Kappa value, *p*_*o*_ is the observed agreement, and *p*_*e*_ is the expected agreement by chance.

## Results

### QS vs NQS classification model (RQ1)

The Gram-Schmidt algorithm was run 10 times using different random seeds on the balanced dataset, yielding the results shown in Table 4. The table reports the number of times each feature ranked in the top five across trials, as well as the average correlation value of each feature. The five strongest features include aEEG-based, EEG-based, and envelope-based features all of which were defined in Table 2.

**Table 4.** Top five strongest features based on their count in the top 5. The table also provides the average correlation of the features with the output across the ten trials.

| Feature | Count | Avg Corr |
| --- | --- | --- |
| aEEG_spectral_flatness | 10 | 0.340 |
| Envelope difference | 10 | 0.310 |
| LOG | 10 | 0.293 |
| D3_AbsMean_Ratio | 10 | 0.287 |
| MYO | 9 | 0.102 |

Regarding the five models trained for QS vs NQS classification, from Table 5, it can be observed that both the k-Nearest Neighbors (KNN) and random forest (RF) models achieved slightly higher values across all metrics we evaluated (accuracy, precision, recall, F1-score, and Kappa).

**Table 5.**
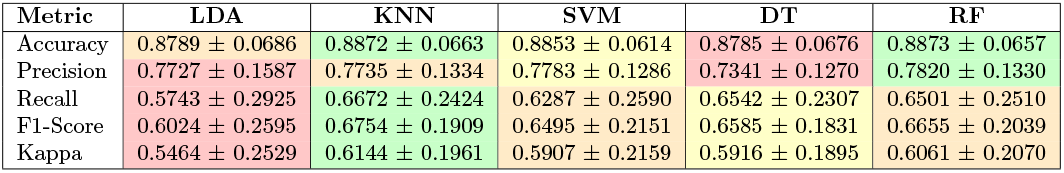
The five trained models in terms of accuracy, precision, recall, f1-score and Kappa when validated using LOSO on 40 neonates.

| Metric | LDA | KNN | SVM | DT | RF |
| --- | --- | --- | --- | --- | --- |
| Accuracy | 0.8789 ± 0.0686 | 0.8872 ± 0.0663 | 0.8853 ± 0.0614 | 0.8785 ± 0.0676 | 0.8873 ± 0.0657 |
| Precision | 0.7727 ± 0.1587 | 0.7735 ± 0.1334 | 0.7783 ± 0.1286 | 0.7341 ± 0.1270 | 0.7820 ± 0.1330 |
| Recall | 0.5743 ± 0.2925 | 0.6672 ± 0.2424 | 0.6287 ± 0.2590 | 0.6542 ± 0.2307 | 0.6501 ± 0.2510 |
| F1-Score | 0.6024 ± 0.2595 | 0.6754 ± 0.1909 | 0.6495 ± 0.2151 | 0.6585 ± 0.1831 | 0.6655 ± 0.2039 |
| Kappa | 0.5464 ± 0.2529 | 0.6144 ± 0.1961 | 0.5907 ± 0.2159 | 0.5916 ± 0.1895 | 0.6061 ± 0.2070 |

To determine whether one model performed statistically better than the other, a Wilcoxon signed-rank test was conducted comparing the RF and KNN models using the Kappa score. The Kappa metric was chosen because it accounts for agreement by chance, providing a more reliable assessment than accuracy for this highly imbalanced classification task (80% NQS, 20% QS). The Wilcoxon test performed on 40 datapoints yielded a test statistic of 325.0000 and a p-value of 0.2591, indicating no significant difference between the performances of the RF and KNN models at the typical significance level (p*<*0.05). For this reason, it was decided to fine-tune only the KNN model, as it was by far the fastest model to train compared to the RF.

Using LOSO cross-validation, the hyperparameter k, which defines the number of neighbours the model considers when classifying a segment, was fine-tuned. At a value of 121 for k, the highest performance in terms of Cohen’s Kappa value was found with a mean value of 0.6159. A detailed table reporting the change in accuracy, F1-score, and Kappa value for each different k value tested is available in Appendix .

The individual Kappa performances for each case in the dataset are presented in Figure 3, while the confusion matrix of each individual case (neonate) can be consulted in S1 Appendix.

**Fig 3.**
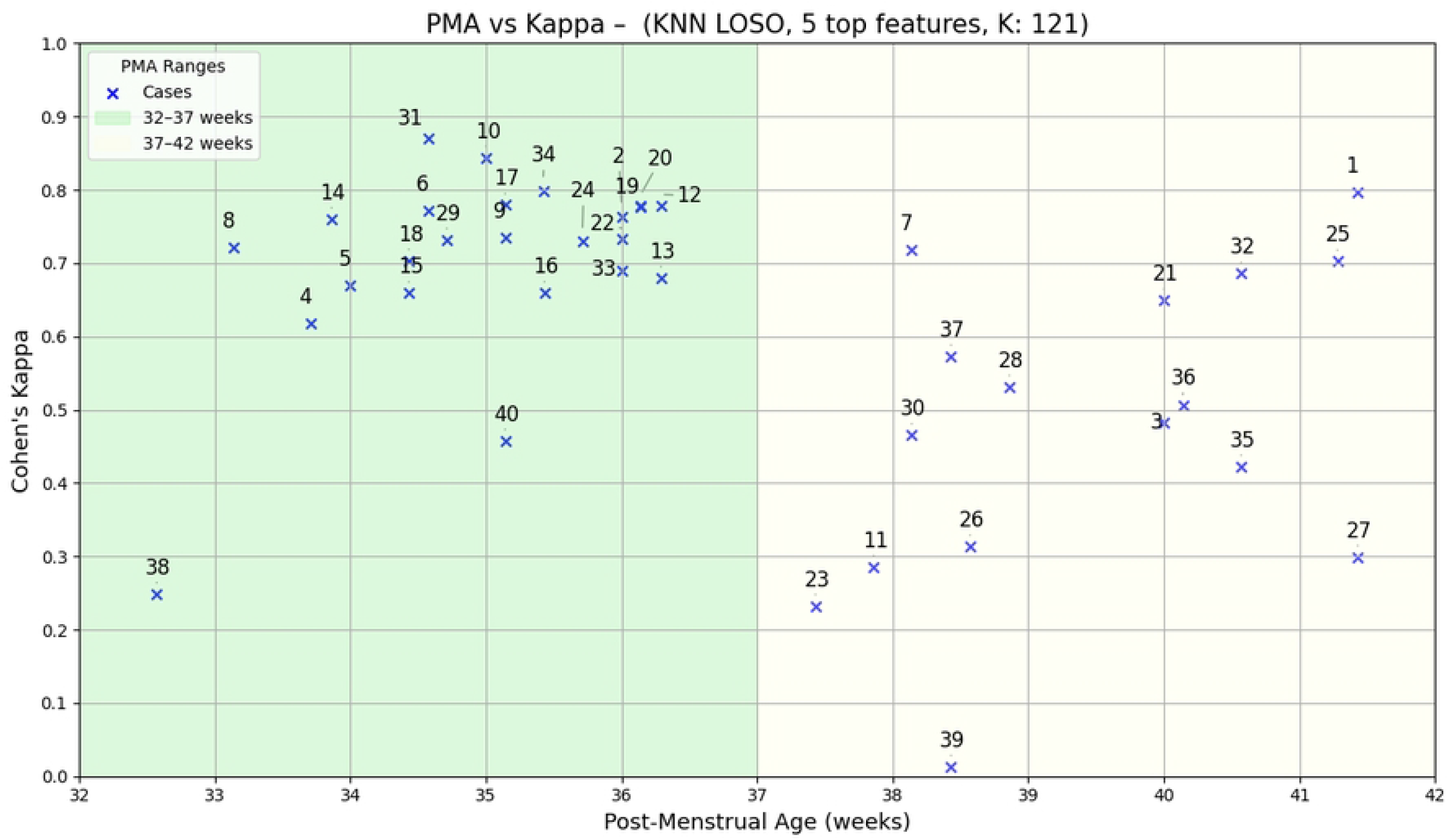
Individual Kappa score on QS versus NQS classification task for each neonate (40 cases) numbered from 1-40 in the dataset. The X-axis displays the PMA of the neonate, while the y-axis shows the performance on the classification task in terms of Kappa.

The results in Figure 3 are also presented in two PMA groups: the preterm (32-37) and near term (37-42) populations. For the preterm group (see green color in Figure 3), the KNN model performed well, with an average Kappa value of 0.69 ± 0.14. Only cases 38 and 40 are outliers to the group and had individual Kappa values of 0.25 and 0.46 respectively. Regarding the term group (see yellow color in Figure 3), the average Kappa value is 0.48 ± 0.21. The results are mixed, ranging from medium to high Kappa values to extremely low values (see case 39 with a Kappa value of 0.01).

### Correlation between QS segments and PMA (RQ2)

The results of the first experiment of RQ2 are shown in Table 6 and Table 7. As can be observed, the values for Set B (see Table 7) are all higher than the correlation values found for Set A (see Table 6). Especially, the ‘Mean lower envelope’ feature calculated on Set B has a positive correlation of 0.602 with the PMA. This suggests that PMA is more correlated with features calculated on QS segments than with those calculated on the original class distribution.

**Table 6.**
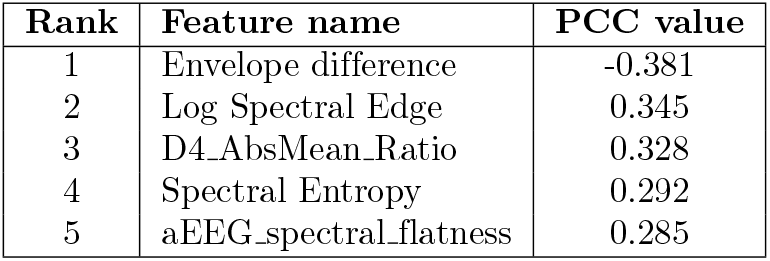
Top 5 features in Set A ranked by Pearson Correlation Coefficient (PCC) with PMA. All correlations are statistically significant with *p <* 0.001.

**Table 7.** Top 5 features in Set B ranked by Pearson Correlation Coefficient (PCC) with PMA. All correlations are statistically significant with *p <* 0.001.

| Rank | Feature name | PCC value |
| --- | --- | --- |
| 1 | Mean lower envelope | 0.602 |
| 2 | Envelope difference | -0.544 |
| 3 | D3_AbsMean_Ratio | -0.501 |
| 4 | MYO | -0.423 |
| 5 | D4_AbsMean_Ratio | 0.396 |

The results of the second experiment, where a model for PMA prediction was trained, further validate this idea. Using set B (containing exclusively QS segments) for training the PLS model yielded a lower MSE and a higher PCC value for all the different values of n components tested (see Figure 4), compared to when using training Set A (having the original 80% NQS, 20% QS distribution). A similar trend for the *R*^2^ metric is visible in Figure S3.

**Fig 4.**
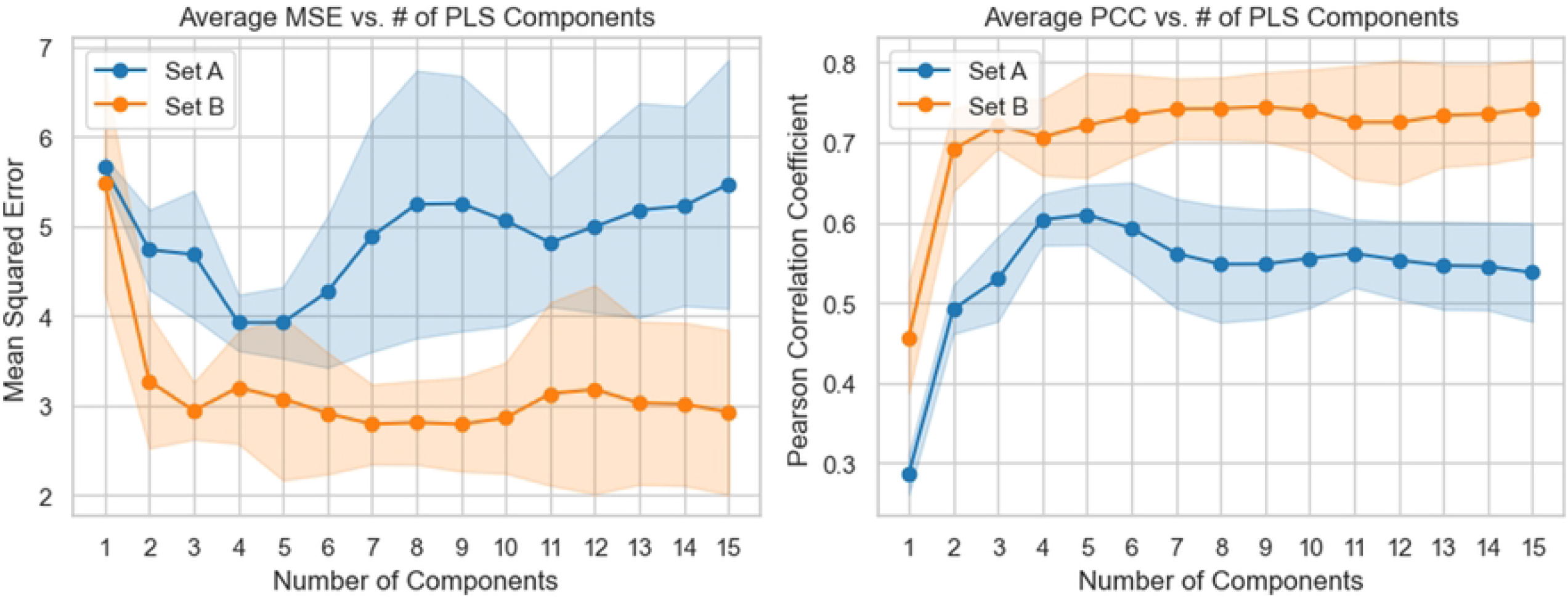
Increasing the value of n components hyperparameter for Set A and Set B. The image displays the average MSE (left plot) and PCC (right plot) value and standard deviation (shaded area) across 10 different random seeds.

Reducing the number of segments selected per neonate provided that from 35 segments, the performances in terms of MSE and *R*^2^ metric stabilized for Set B, as can be observed in Figure 5.

**Fig 5.**
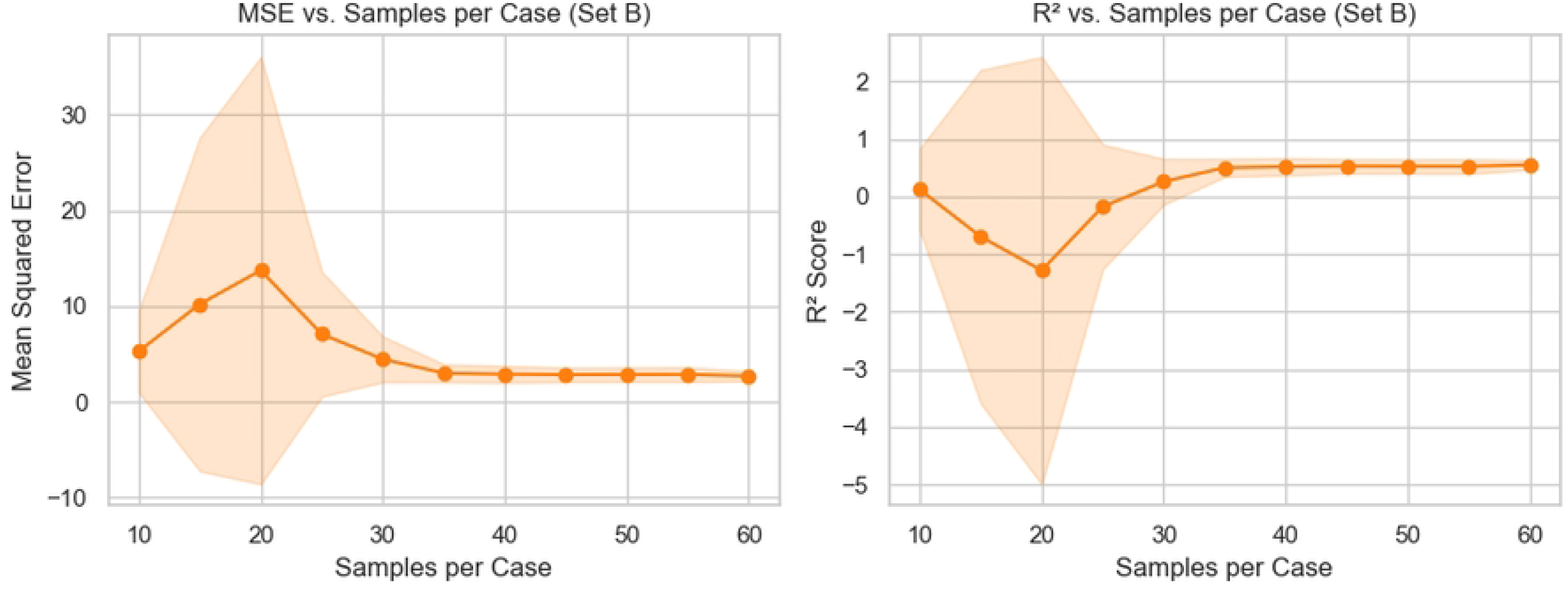
Effect of increasing the number of QS segments selected per neonate to constitute training set B on MSE and *R*^2^ metric

Finally, using this reduced number of selected QS segments per neonate (N = 35) a model was trained for PMA prediction using LOSO cross-validation (39 neonates for training, 1 for testing). Here ‘n components’ hyperparameter was set to nine, as from Figure 4 it could be seen that this was the lowest value for ‘n components’ achieving the highest performances both in MSE and PCC. The results of this model are visible in Figure 6. The average PMA predicted and its standard deviation are both visible on the image, compared with the True PMA of each neonate. This average prediction is calculated based on the 35 PMA predictions that the model makes for each neonate (one for each segment provided). The final model performances in terms of PCC, MSE and average error across the 40 neonates are: 0.91, 1.33 weeks and 0.88 weeks respectively.

**Fig 6.**
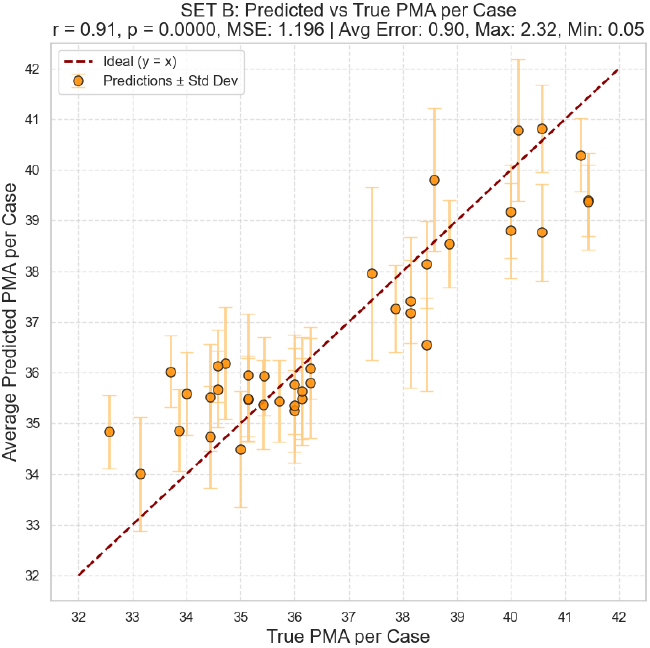
Partial Least Square model average PMA predictions compared with True PMA of 40 different neonates.

In order to evaluate the feasibility of a fully automated, end-to-end pipeline for brain maturation assessment, a third dataset, Set C, was created. In this setup, QS segments were not annotated by the clinician, but instead predicted by the KNN model trained previously. For each neonate, 35 predicted QS segments were randomly selected and used as input to the same PLS model for PMA prediction.

However, for case 39, the KNN model did not predict a sufficient number of QS segments to allow for the selection of 35 valid samples. As a result, this neonate was excluded from the Set C-based PMA evaluation. The final model performance metrics were therefore computed over the remaining 39 neonates. The results of this model PMA prediction on 39 neonates are reported in Figure 7. The model achieved a MSE of 1.77 weeks, an average error of 1.08 weeks, and a PCC of 0.86.

**Fig 7.**
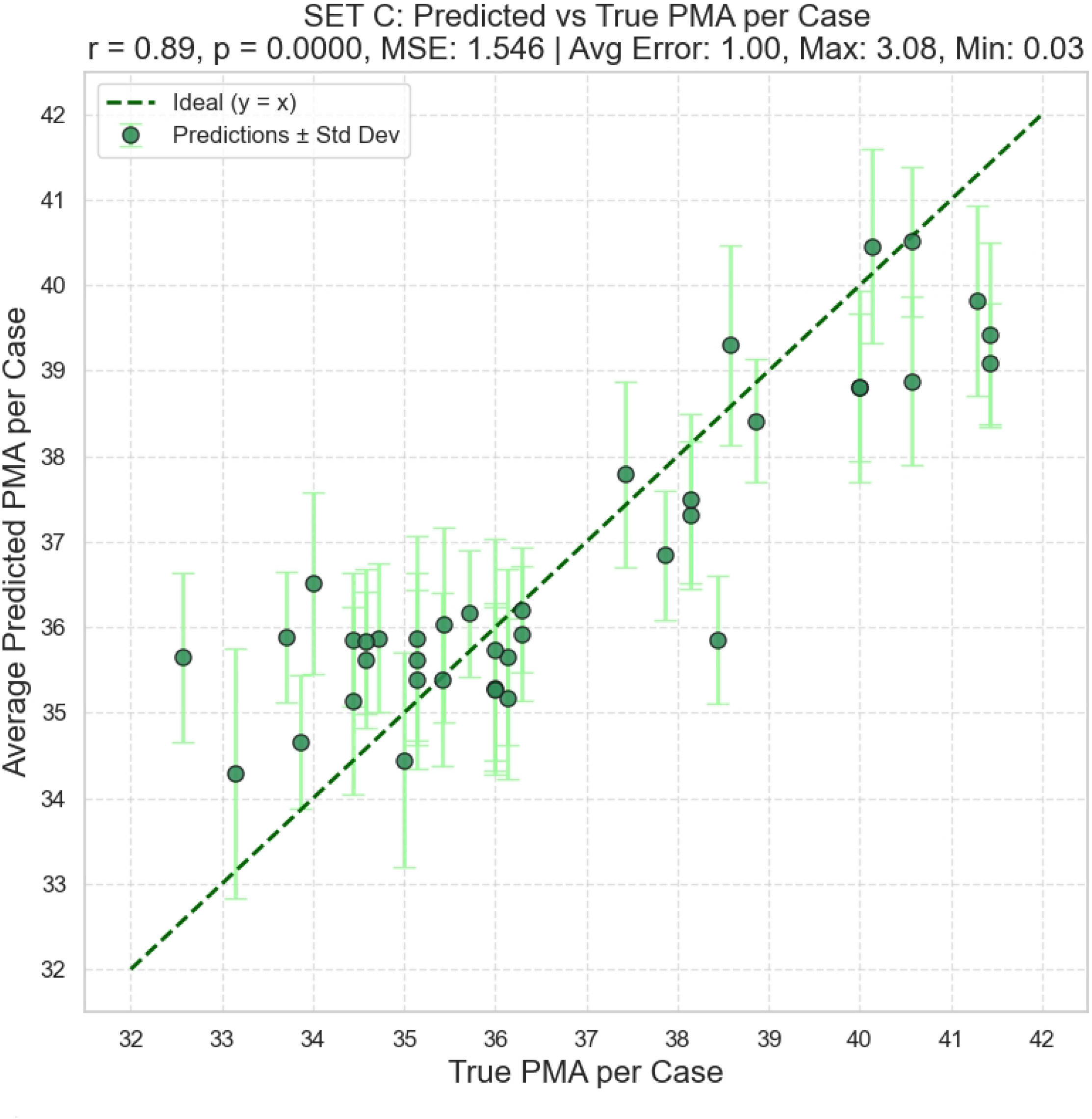
Partial Least Square model average PMA predictions compared with True PMA of 39 different neonates.

## Discussion

Unlike prior single-channel studies that either down-select from PSG or focus on term infants [35, 40, 41, 43, 44], our dataset comprises long aEEG-standard recordings across 32–42 weeks PMA, directly targeting clinical reality.

While we separated the neonates into two groups, the preterms *<* 37 weeks on one side, and the near-terms *>* 37 weeks on the other side, the results show that ideally a third category should be created (lower than 33 weeks) for the youngest neonate: case 38 with 32.56 weeks PMA. This neonate, which stood out from the rest of the preterm group with a low Kappa value of 0.25, also differed from the other neonates at 33 weeks of PMA during visual inspection of the aEEG trend. For illustration, Figure S1 presents the aEEG trace of the youngest neonate in our cohort (32.56 weeks PMA), where the quiet sleep (QS) period is highlighted in green and the upper and lower signal envelopes are depicted in red; and Figure S2 shows the recording of the second youngest participant (33.14 weeks PMA),

As observed in Figure S1, the envelope difference for the youngest neonate remains large over the whole recording, making the periods of QS less distinguishable from the rest of the aEEG trace. In contrast, for the second neonate (see Figure S2), portions of the aEEG trend envelope are narrower (i.e., smaller envelope difference) before and after each QS period, which makes the periods of QS more clearly identifiable.

The other outlier in the preterm group is case 40, which had a Kappa value of 0.46. For this case, the medical record of this neonate reported that the neonate received caffeine treatment during the recording, affecting the duration of its sleep-wake cycles (SWC).

Direct numerical comparison is limited by different sensors (aEEG vs PSG), taxonomy (QS/NQS vs multi-stage), cohorts (preterm vs term), and validation schemes (LOSO vs k-fold); nevertheless, our *κ* values are within the range of prior single-channel reports [35, 40, 41] while addressing the preterm aEEG gap.

The duration of this neonate’s SWC was on average 11.45 minutes compared to a 19.09 ± 5.94 minutes average for the whole preterm group. The z-score analysis of this case also reported a z-value of -2.56 with a p-value of 0.01, which statistically confirms that this case is an outlier to the group in terms of SWC durations.

We believe that these shorter SWC durations may have affected some of the features used by the KNN model to perform the QS/NQS classification, which can explain the lower Kappa value found for this case.

Regarding the term group, starting at 37 weeks, the classifier showed mixed performance. Middle to high range Kappa values were reported for the older cases, but extremely low values were observed in some cases at 38.5 weeks of PMA (see case 39). This suggests that the data in this group is very heterogeneous as two cases with similar PMA (see case 37 and 39) performed very differently in terms of Kappa value.

Regarding comparison with other literature, Ansari et al. had a Kappa value of 0.75 on the validation set, but as they used a smaller database (only 26 neonates), a different data acquisition system (no aEEG data), and did not report the exact PMA of the neonates on which they validated their model, it is difficult to compare our model with the performances they reported.

The answer to our first research question is therefore that while it was possible to train a model performing well across the 32-37 weeks PMA range, it was not possible to train a single model that generalizes well for all the PMA categories present in NICUs. More data at a PMA range of 37-38 weeks is required to understand how the data evolves with increasing PMA.

Regarding the second research question, a higher correlation value was found when computing the PCC of each individual feature with PMA using QS segments only. Both the ‘Mean lower envelope’ and the ‘Envelope difference’ correlated strongly with the PMA of the neonate. Interestingly, this coincides with two parameters of the Burdjalov scoring system: the lower border (LB) and aEEG signal bandwidth (B).

Using exclusively QS segments for training a Partial Least Squares (PLS) model to predict PMA resulted in improved performance, as indicated by the higher PCC, higher *R*^2^, and lower MSE metrics found, compared to the model trained on the original data distribution (80% NQS / 20% QS).

Our PLS model trained using the 35 expert-labeled QS segments from each case had an average error of 0.88 weeks, a mean square error of 1.33 weeks, and a maximum error of 2.70 weeks. This demonstrates the potential of such an automated solution for estimating brain maturation as an objective alternative to the Burdjalov scoring system, especially considering that a study by Stevenson et al. [49] reported a random error of 1.8 weeks among expert clinicians using aEEG for brain maturation prediction.

In addition, since our model requires only 35 QS segments (each 30 seconds long) for training, the duration of the aEEG recording for each neonate could potentially be reduced. On average, a recording of 24 hours contains well over 100 segments annotated as QS. Therefore, in an exploratory approach, a small proof was conducted where only the first 12 hours of data from each case of the database were kept. Randomly selecting the 35 QS segments in these 12 hours of data provided similar results to those presented in Section . Only the average error had a slight increase from 0.88 to 0.96 weeks.

Finally, in an attempt to test a fully integrated pipeline, segments predicted as QS by the KNN model were used to train a PLS model to predict PMA (Set C). This PLS model achieved a final average error of 1.08 weeks, a mean squared error of 1.77 weeks, and a maximum error of 3.02 weeks. While these results are not fully comparable to those obtained using manual expert-made annotations—because the model failed to predict enough QS segments for one neonate, resulting in evaluation on one fewer subject—they still demonstrate that a fully integrated pipeline holds promise as a solution that eliminates human variability in PMA prediction.

Even though particular effort was made to help the clinician annotate the dataset accurately, we recognize that some segments might still be misclassified because of artefacts and interrupted SWC. This could penalize both the training and validation of the classifier model, as mislabeled segments may confuse the model during training and lead to inaccurate evaluations during validation.

Moreover, a key limitation of this study is that the annotation of QS segments was performed by a single expert clinician. Although specific guidelines and visual aids were provided to enhance consistency, the lack of inter-rater validation may introduce subjectivity and bias in the reference labels.

Additionally, our dataset contained an unbalanced amount of cases around 33-36 weeks of PMA. This is a condition that might have biased the model to be stronger around these PMA weeks. However, training the model using a smaller but more balanced dataset did not improve the model’s abilities to generalize well over all PMA categories.

Finally, due to computational constraints, some more computationally demanding models such as a non-linear kernel support vector machine or neural network could not be explored. Also, the choice for keeping only the top five features is a limitation of the study. While it allows for a fast training of the models, exploiting more features could potentially leverage powerful feature combinations leading to higher performances.

For future work, it could be interesting to test the found KNN model on more data recorded from the 33-37 category to see if it further generalizes for this group.

We also believe it would be valuable to identify features that have strong predictive value for classifying QS/NQS, but that are not correlated with the PMA of the neonates. This would ensure that these features are not specific to sleep detection for certain PMA categories, and potentially allow for better generalization across all categories.

Regarding the second research question, the performances of the PMA predictor model should be compared with a group of expert clinicians predicting the age of the same neonates included in this study using the Burdjalov scoring system. This would allow for a fair comparison of the presented PMA predictor model with the current clinical reality.

## Conclusion

This study provides, to our knowledge, the first QS/NQS approach validated on preterm aEEG-standard data, aligning method and clinical context.

The development of a model for QS/NQS classification that performs reliably across a broad range of PMAs is currently hindered by the substantial differences in aEEG patterns between neurologically healthy preterm neonates aged 32–37 weeks and those from 37 weeks onward. Still, this study presented a KNN solution that achieved a mean Kappa value of 0.69 ± 0.14 for neonates between 32 and 37 weeks working on a single-channel EEG signal, recorded using aEEG standards.

Features calculated on QS segments were more correlated with PMA than features calculated on an equally sized set containing the original (80% NQS, 20% QS) class distribution. The filtering of QS segments to train a model for PMA prediction also yielded a higher PCC and lower MSE metric compared to training with a set following the original (80% NQS, 20% QS) class distribution.

Finally, the model trained for PMA prediction had an average error of 0.88 weeks. These findings highlight the potential of automated PMA prediction approaches through (automated) QS classification as a more precise and objective alternative to current subjective methods used for mild brain dysfunction assessment in preterm infants.

## Acknowledgments

We would like to express our gratitude to the healthcare professionals Gisel Montoya Aguirre, Susana Larrosa Capaces, Montse Roselló Foguet, and Vanesa Rius Costa from the Clinical Neurophysiology Service at HUSJR for their valuable collaboration. We also thank Ismael Ávila and Agnès Rigo Vidal for their support in the development of the clinical research plan.

## Supporting information

### Supplementary File

**Fig S1.**
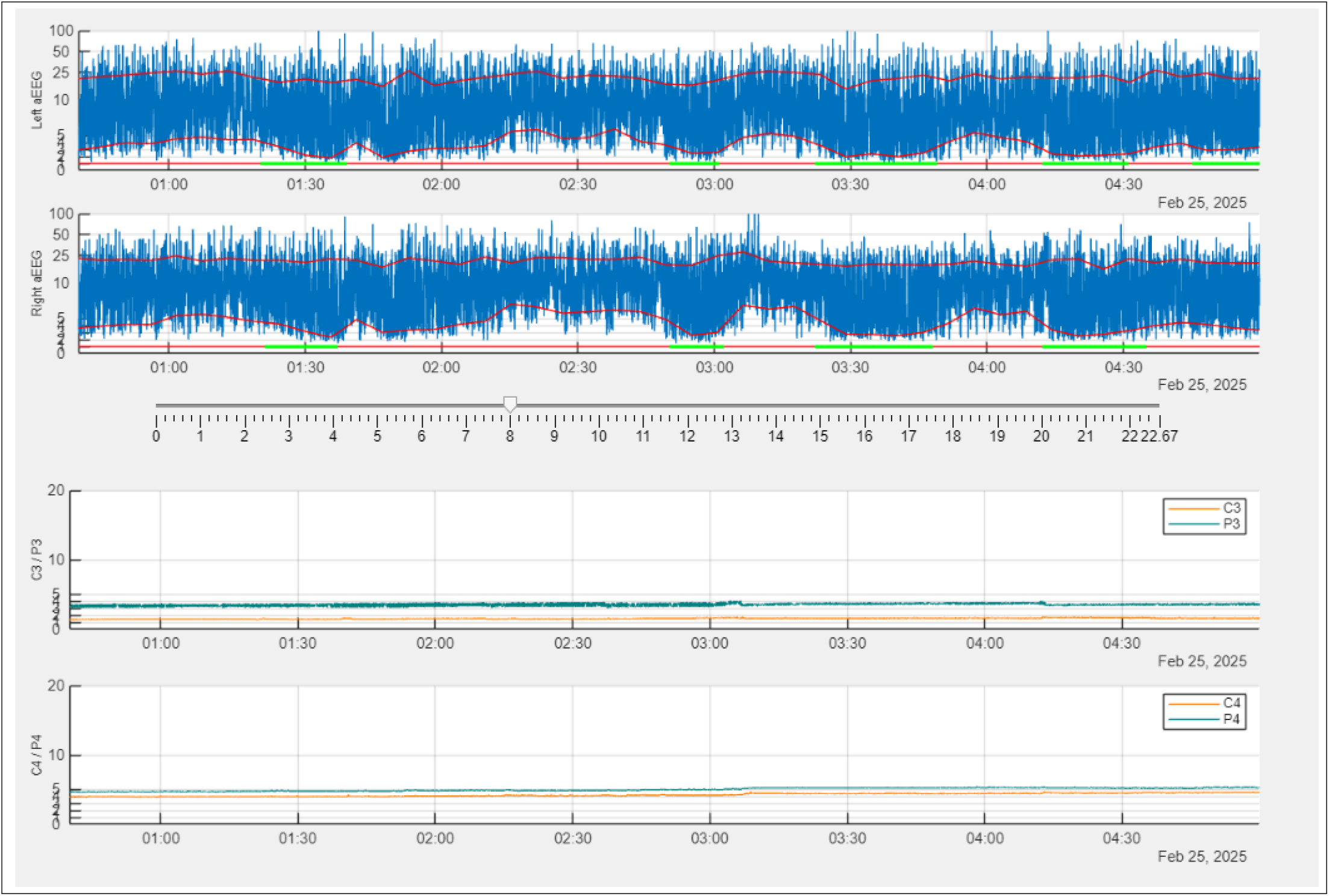
Screenshot of case 38, aEEG view of the youngest neonate in our cohort at 32.56 weeks. Sleep-wake cycle annotated in green on aEEG signal together with upper and lower envelopes (red lines) of the signal showing general trend of the aEEG.

**Fig S2.**
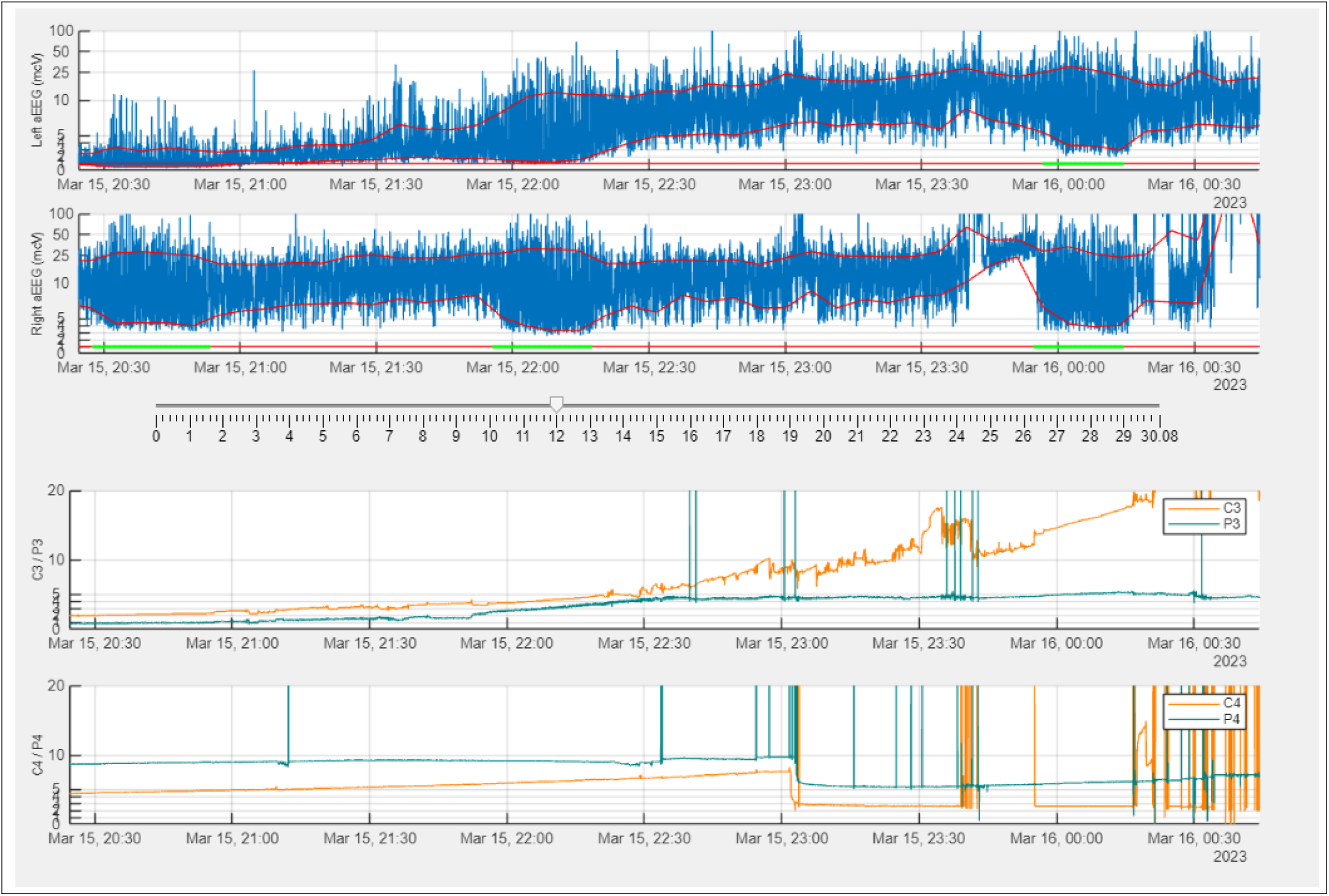
Screenshot of case 8, aEEG view of the second youngest neonate in our cohort at 33.14 weeks. Sleep-wake cycle annotated in green on aEEG signal together with upper and lower envelopes (red lines) of the signal showing general trend of the aEEG.

**Fig S3.**
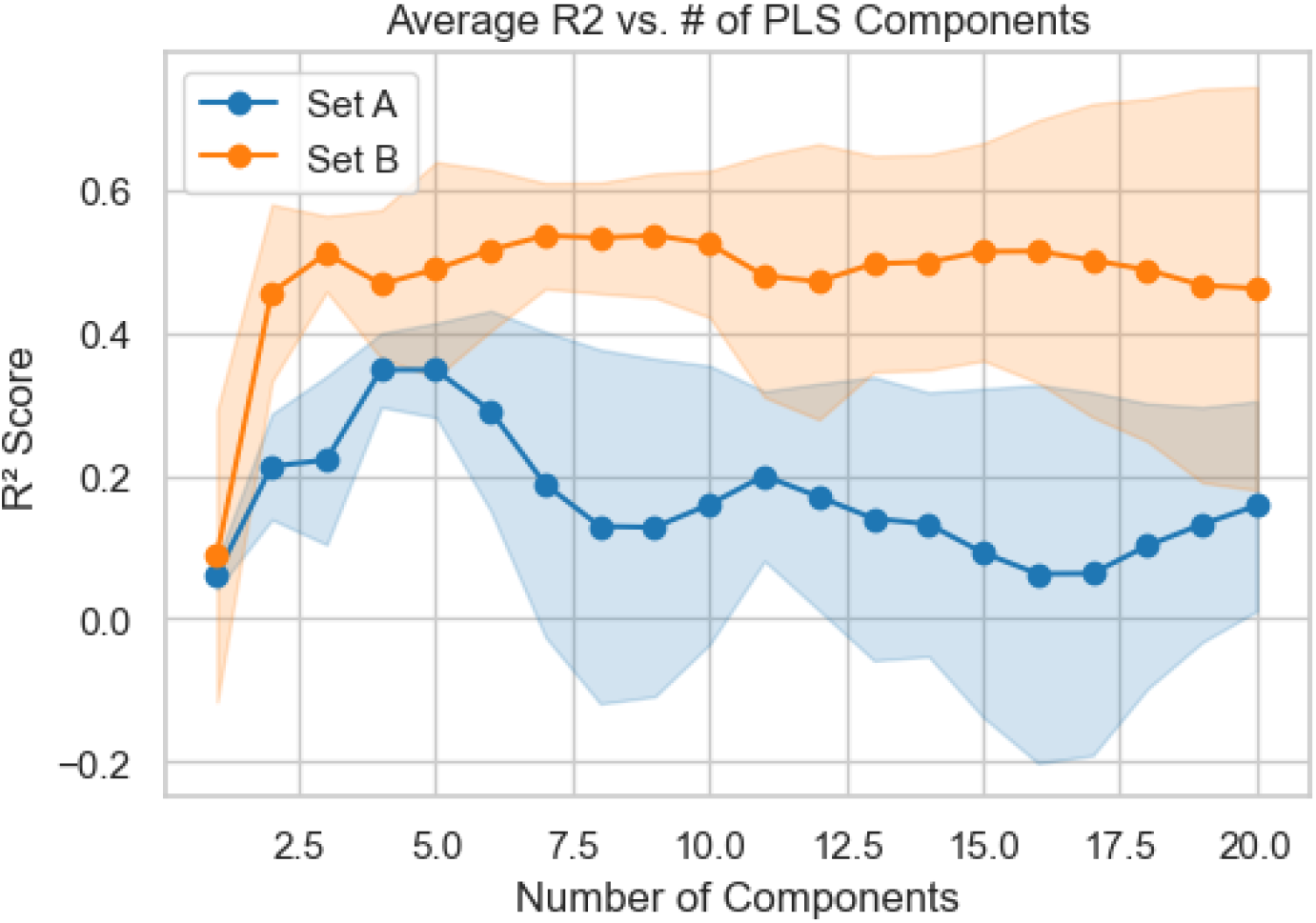
*R*^2^ average values and standard deviation for training using set A and B across 10 different random seeds.

**Table S1.**
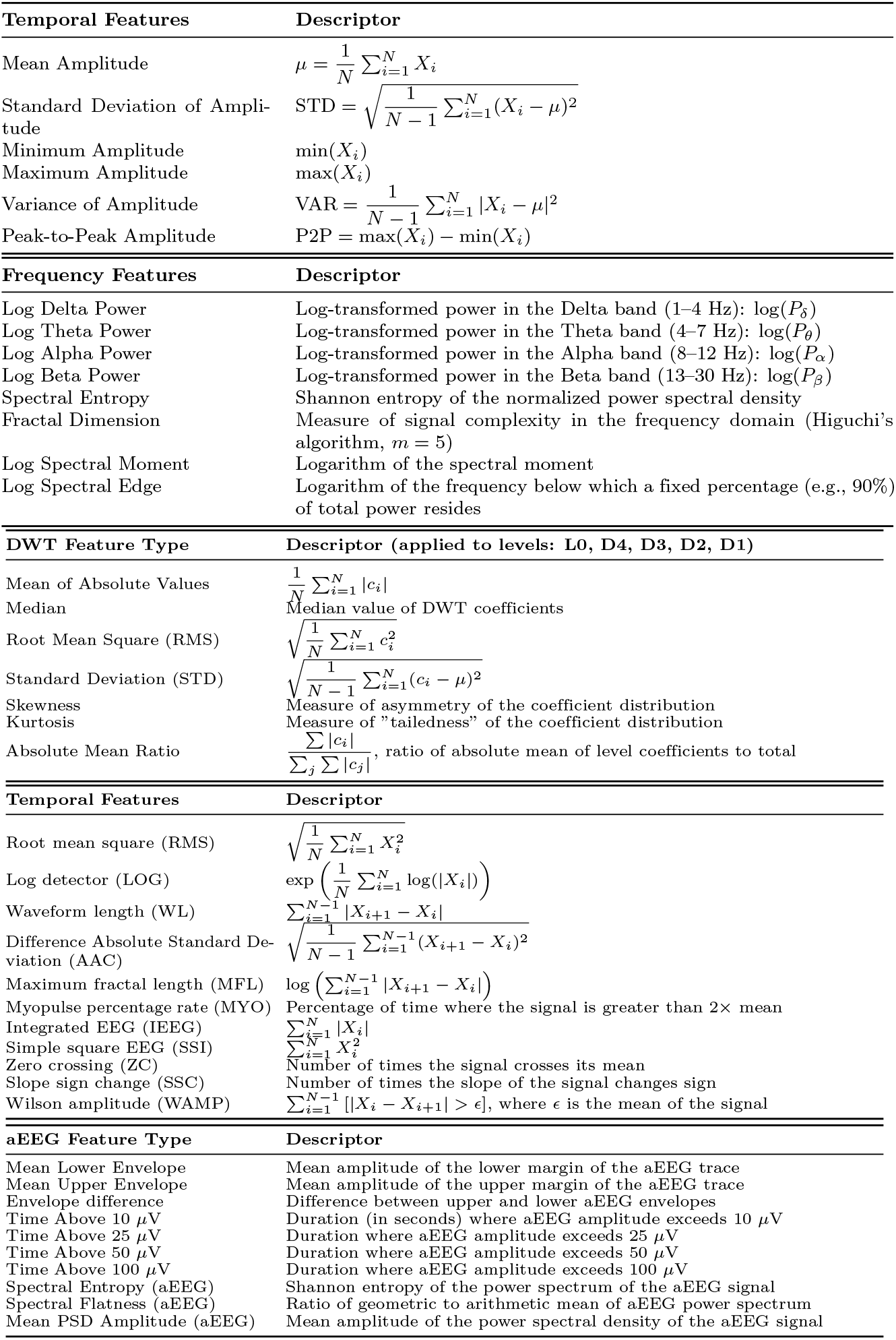
Comprehensive list of extracted features used in this study, including EEG-based features in the temporal domain, EEG-based features in the frequency domain of the 30 s segments, EEG-based Discrete Wavelet Transform (DWT) features, EMG-inspired features computed on 30 s EEG segments, and amplitude-integrated EEG-based features.

**Table S2.** Performance Metrics, K-value for KNN model.

| k | Mean Accuracy | Mean Kappa | Mean F1-score |
| --- | --- | --- | --- |
| 1 | 0.8376 | 0.4826 | 0.5786 |
| 6 | 0.8758 | 0.5575 | 0.6233 |
| 11 | 0.8818 | 0.6015 | 0.6666 |
| 16 | 0.8841 | 0.5987 | 0.6608 |
| 21 | 0.8847 | 0.6096 | 0.6724 |
| 26 | 0.8852 | 0.6046 | 0.6663 |
| 31 | 0.8858 | 0.6122 | 0.6743 |
| 36 | 0.8861 | 0.6092 | 0.6706 |
| 41 | 0.8866 | 0.6158 | 0.6775 |
| 46 | 0.8871 | 0.6134 | 0.6744 |
| 51 | 0.8868 | 0.6157 | 0.6772 |
| 56 | 0.8870 | 0.6136 | 0.6746 |
| 61 | 0.8868 | 0.6155 | 0.6769 |
| 66 | 0.8870 | 0.6136 | 0.6746 |
| 71 | 0.8870 | 0.6157 | 0.6770 |
| 76 | 0.8871 | 0.6140 | 0.6748 |
| 81 | 0.8869 | 0.6154 | 0.6766 |
| 86 | 0.8870 | 0.6140 | 0.6749 |
| 91 | 0.8871 | **0.6159** | 0.6770 |
| 96 | 0.8873 | 0.6148 | 0.6757 |
| 101 | 0.8869 | 0.6152 | 0.6765 |
| 106 | 0.8870 | 0.6141 | 0.6751 |
| 111 | 0.8871 | 0.6155 | 0.6767 |
| 116 | 0.8872 | 0.6144 | 0.6752 |
| 121 | 0.8872 | **0.6159** | 0.6770 |
| 126 | 0.8870 | 0.6141 | 0.6751 |
| 131 | 0.8868 | 0.6146 | 0.6758 |
| 136 | 0.8872 | 0.6144 | 0.6752 |
| 141 | 0.8871 | 0.6154 | 0.6764 |
| 146 | 0.8872 | 0.6144 | 0.6752 |
| 151 | 0.8872 | 0.6155 | 0.6765 |
| 156 | 0.8872 | 0.6140 | 0.6748 |
| 161 | 0.8870 | 0.6142 | 0.6753 |
| 166 | 0.8871 | 0.6139 | 0.6748 |
| 171 | 0.8872 | 0.6151 | 0.6760 |

### S1 Appendix Confusion Matrices of KNN model using five features and K:121

|  | Predicted 0 | Predicted 1 |
| --- | --- | --- |
| Actual 0 | 2035 | 137 |
| Actual 1 | 61 | 531 |

Case: 2, PMA: 36.0, Kappa: 0.764
|  | Predicted 0 | Predicted 1 |
| --- | --- | --- |
| Actual 0 | 1682 | 40 |
| Actual 1 | 111 | 314 |

Case: 3, PMA: 40.0, Kappa: 0.483
|  | Predicted 0 | Predicted 1 |
| --- | --- | --- |
| Actual 0 | 3938 | 20 |
| Actual 1 | 658 | 407 |

Case: 4, PMA: 33.71, Kappa: 0.619
|  | Predicted 0 | Predicted 1 |
| --- | --- | --- |
| Actual 0 | 4350 | 61 |
| Actual 1 | 467 | 540 |

Case: 5, PMA: 34.0, Kappa: 0.669
|  | Predicted 0 | Predicted 1 |
| --- | --- | --- |
| Actual 0 | 2239 | 299 |
| Actual 1 | 93 | 582 |

Case: 6, PMA: 34.57, Kappa: 0.772
|  | Predicted 0 | Predicted 1 |
| --- | --- | --- |
| Actual 0 | 3228 | 157 |
| Actual 1 | 159 | 711 |

Case: 7, PMA: 38.14, Kappa: 0.719
|  | Predicted 0 | Predicted 1 |
| --- | --- | --- |
| Actual 0 | 2782 | 116 |
| Actual 1 | 53 | 251 |

**Case: 8, PMA: 33.14, Kappa: 0.722**
|  | Predicted 0 | Predicted 1 |
| --- | --- | --- |
| Actual 0 | 847 | 103 |
| Actual 1 | 55 | 330 |

**Case: 9, PMA: 35.14, Kappa: 0.735**
|  | Predicted 0 | Predicted 1 |
| --- | --- | --- |
| Actual 0 | 648 | 55 |
| Actual 1 | 65 | 273 |

**Case: 10, PMA: 35.0, Kappa: 0.844**
|  | Predicted 0 | Predicted 1 |
| --- | --- | --- |
| Actual 0 | 1574 | 75 |
| Actual 1 | 39 | 414 |

**Case: 11, PMA: 37.86, Kappa: 0.286**
|  | Predicted 0 | Predicted 1 |
| --- | --- | --- |
| Actual 0 | 2959 | 5 |
| Actual 1 | 478 | 117 |

**Case: 12, PMA: 36.29, Kappa: 0.779**
|  | Predicted 0 | Predicted 1 |
| --- | --- | --- |
| Actual 0 | 1493 | 16 |
| Actual 1 | 91 | 232 |

**Case: 13, PMA: 36.29, Kappa: 0.680**
|  | Predicted 0 | Predicted 1 |
| --- | --- | --- |
| Actual 0 | 1996 | 69 |
| Actual 1 | 122 | 252 |

**Case: 14, PMA: 33.86, Kappa: 0.760**
|  | Predicted 0 | Predicted 1 |
| --- | --- | --- |
| Actual 0 | 1879 | 80 |
| Actual 1 | 109 | 398 |

**Case: 15, PMA: 34.43, Kappa: 0.659**
|  | Predicted 0 | Predicted 1 |
| --- | --- | --- |
| Actual 0 | 740 | 82 |
| Actual 1 | 37 | 163 |

**Case: 16, PMA: 35.43, Kappa: 0.659**
|  | Predicted 0 | Predicted 1 |
| --- | --- | --- |
| Actual 0 | 1449 | 29 |
| Actual 1 | 213 | 331 |

**Case: 17, PMA: 35.14, Kappa: 0.779**
|  | Predicted 0 | Predicted 1 |
| --- | --- | --- |
| Actual 0 | 1065 | 68 |
| Actual 1 | 31 | 231 |

**Case: 18, PMA: 34.43, Kappa: 0.703**
|  | Predicted 0 | Predicted 1 |
| --- | --- | --- |
| Actual 0 | 1536 | 99 |
| Actual 1 | 78 | 277 |

**Case: 19, PMA: 36.14, Kappa: 0.777**
|  | Predicted 0 | Predicted 1 |
| --- | --- | --- |
| Actual 0 | 1353 | 80 |
| Actual 1 | 45 | 286 |

**Case: 20, PMA: 36.14, Kappa: 0.778**
|  | Predicted 0 | Predicted 1 |
| --- | --- | --- |
| Actual 0 | 1975 | 80 |
| Actual 1 | 86 | 374 |

**Case: 21, PMA: 40.0, Kappa: 0.649**
|  | Predicted 0 | Predicted 1 |
| --- | --- | --- |
| Actual 0 | 2007 | 247 |
| Actual 1 | 81 | 428 |

**Case: 22, PMA: 35.0, Kappa: 0.733**
|  | Predicted 0 | Predicted 1 |
| --- | --- | --- |
| Actual 0 | 1794 | 155 |
| Actual 1 | 43 | 359 |

**Case: 23, PMA: 37.43, Kappa: 0.233**
|  | Predicted 0 | Predicted 1 |
| --- | --- | --- |
| Actual 0 | 2460 | 40 |
| Actual 1 | 171 | 38 |

**Case: 24, PMA: 35.71, Kappa: 0.729**
|  | Predicted 0 | Predicted 1 |
| --- | --- | --- |
| Actual 0 | 2120 | 134 |
| Actual 1 | 105 | 430 |

**Case: 25, PMA: 40.14, Kappa: 0.703**
|  | Predicted 0 | Predicted 1 |
| --- | --- | --- |
| Actual 0 | 968 | 77 |
| Actual 1 | 47 | 199 |

**Case: 26, PMA: 37.0, Kappa: 0.314**
|  | Predicted 0 | Predicted 1 |
| --- | --- | --- |
| Actual 0 | 1920 | 103 |
| Actual 1 | 452 | 201 |

**Case: 27, PMA: 41.43, Kappa: 0.299**
|  | Predicted 0 | Predicted 1 |
| --- | --- | --- |
| Actual 0 | 2041 | 26 |
| Actual 1 | 348 | 103 |

**Case: 28, PMA: 38.86, Kappa: 0.531**
|  | Predicted 0 | Predicted 1 |
| --- | --- | --- |
| Actual 0 | 1597 | 95 |
| Actual 1 | 157 | 192 |

**Case: 29, PMA: 34.71, Kappa: 0.732**
|  | Predicted 0 | Predicted 1 |
| --- | --- | --- |
| Actual 0 | 1936 | 39 |
| Actual 1 | 132 | 292 |

**Case: 30, PMA: 38.14, Kappa: 0.465**
|  | Predicted 0 | Predicted 1 |
| --- | --- | --- |
| Actual 0 | 2663 | 97 |
| Actual 1 | 252 | 192 |

**Case: 31, PMA: 34.57, Kappa: 0.870**
|  | Predicted 0 | Predicted 1 |
| --- | --- | --- |
| Actual 0 | 2211 | 68 |
| Actual 1 | 46 | 486 |

**Case: 32, PMA: 40.57, Kappa: 0.686**
|  | Predicted 0 | Predicted 1 |
| --- | --- | --- |
| Actual 0 | 1702 | 95 |
| Actual 1 | 124 | 322 |

**Case: 33, PMA: 36.0, Kappa: 0.689**
|  | Predicted 0 | Predicted 1 |
| --- | --- | --- |
| Actual 0 | 3532 | 153 |
| Actual 1 | 475 | 1048 |

**Case: 34, PMA: 35.42, Kappa: 0.799**
|  | Predicted 0 | Predicted 1 |
| --- | --- | --- |
| Actual 0 | 2102 | 78 |
| Actual 1 | 55 | 323 |

**Case: 35, PMA: 40.57, Kappa: 0.423**
|  | Predicted 0 | Predicted 1 |
| --- | --- | --- |
| Actual 0 | 2722 | 44 |
| Actual 1 | 522 | 280 |

**Case: 36, PMA: 41.14, Kappa: 0.506**
|  | Predicted 0 | Predicted 1 |
| --- | --- | --- |
| Actual 0 | 1354 | 536 |
| Actual 1 | 85 | 649 |

**Case: 37, PMA: 38.29, Kappa: 0.573**
|  | Predicted 0 | Predicted 1 |
| --- | --- | --- |
| Actual 0 | 869 | 41 |
| Actual 1 | 148 | 189 |

**Case: 38, PMA: 32.57, Kappa: 0.249**
|  | Predicted 0 | Predicted 1 |
| --- | --- | --- |
| Actual 0 | 1141 | 1097 |
| Actual 1 | 20 | 460 |

**Case: 39, PMA: 38.43, Kappa: 0.013**
|  | Predicted 0 | Predicted 1 |
| --- | --- | --- |
| Actual 0 | 2231 | 0 |
| Actual 1 | 779 | 7 |

Case: 40, PMA: 35.14, Kappa: 0.458
|  | Predicted 0 | Predicted 1 |
| --- | --- | --- |
| Actual 0 | 2515 | 42 |
| Actual 1 | 156 | 96 |

